# Chemotherapeutic dose scheduling based on tumor growth rates: *the case for low dose metronomic high entropy therapies*

**DOI:** 10.1101/166058

**Authors:** Jeffrey West, Paul K. Newton

**Keywords:** metronomic chemotherapy, evolutionary game theory, prisoner’s dilemma, computational methods, tumor progression

## Abstract

We extend classical tumor regression models, such as the Norton-Simon hypothesis, from instantaneous regression rates (i.e. the derivative) to the cumulative effect (i.e. the integral) over one (or many) cycles of chemotherapy. To achieve this end, we use a stochastic Moran process model of tumor cell kinetics, coupled with a prisoner’s dilemma game-theoretic cell-cell interaction model to design chemotherapeutic strategies tailored to different tumor growth characteristics. Using the Shannon entropy as a novel tool to quantify the success of dosing strategies, we contrast maximum tolerated dose (MTD) strategies as compared with low dose, high density metronomic strategies (LDM) for tumors with different growth rates. Our results show that LDM strategies can outperform MTD strategies in total tumor cell reduction (TCR). The advantage is magnified for fast growing tumors that thrive on long periods of unhindered growth without chemotherapy drugs present and is not evident after a single cycle of chemotherapy, but grows after each subsequent cycle of repeated chemotherapy. The model supports the concept of designing different chemotherapeutic schedules for tumors with different growth rates and develops quantitative tools to optimize these schedules for maintaining low volume tumors. The evolutionary model we introduce in this paper is compared with regression data from murine models and shown to be in good agreement.

**Major Findings:** Model simulations show that metronomic (low dose, high density) therapies can outperform maximum tolerated dose (high dose, low density) therapies. This is due to the fact that tumor cell reduction is more sensitive to changes in dose density than changes in dose concentration, especially for faster growing tumors. This effect is negligible after a single cycle of chemotherapy, but magnified after many cycles. The model also allows for novel chemotherapeutic schedules and quantifies their performance according to tumor growth rate.

## Quick Guide to Equations and Assumptions

### Assumptions of the model

1. The model is a computational one, driven by a stochastic Moran (birth-death) process with a finite cell population, in which birth-death rates are functions of cell fitness.
2. Two classes of cells (healthy, cancer) compete against each other at each birth-death event, with fitness (*f_H_*, *f_C_*) calculated according to the payoff matrix associated with the prisoners dilemma evolutionary zero-sum game.
3. Chemotherapy is a selective agent that alters the selection pressure (*w_H_*, *w_C_*) of each cell population, with two parameters: dose concentration (*c*) and dose density (*d*).

### Key equations

In a Moran finite-population, birth-death process there are *i* cancer cells in a population of *N* total cells (where the number of healthy cells is denoted *N − i*). At each time step in the stochastic evolutionary population dynamics model, a single cell is chosen for birth and a separate single cell is chosen for death. A tumor grows by increasing the number of cancer cells from *i* to *i* + 1 in any given time step. The probability that a healthy cell interacts with another healthy cell is given by (*N − i −* 1)*/*(*N −* 1), whereas the probability that a healthy cell interacts with a cancer cell is *i/*(*N −* 1). The probability that a cancer cell interacts with a healthy cell is (*N − i*)*/*(*N −* 1), whereas the probability that a cancer cell interacts with another cancer cell is (*i −* 1)*/*(*N −* 1). These probabilities, known as the Moran process, can be extended to include a fitness landscape where natural selection can play out over many cell division timescales.

The probabilities outlined above are weighted by the “payoff” in order to determine the fitness function for each subpopulation: healthy (*f_H_*) and cancer (*f_C_*), below. The payoff values (*a*, *b*, *c*, *d*) are associated with the prisoner’s dilemma evolutionary game (1, 2). The prisoner’s dilemma is defined by the payoff inequalities such that *c > a > d > b*, but here we assume the relatively standard (but not unique) values of *a* = 3, *b* = 0, *c* = 5, and *d* = 1.

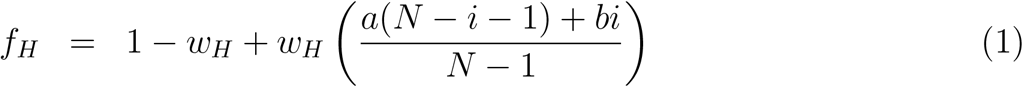

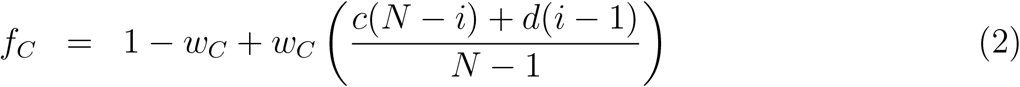

Here, (*w_H_*, *w_C_*) are “selection strength” parameters, 0 *≤ w_H_ ≤* 1, 0 *≤ w_C_ ≤* 1, that measure the strength of selection pressure on each of the population of cells. If *w_H_* = 0, there is no natural selection acting on the healthy cell population, and the dynamics is driven purely by the Moran process (i.e. random drift). When *w_H_* = 1, the selection pressure on the healthy cell population is strongest, and the prisoners dilemma payoff matrix has maximum effect.

From these formulas, we can define the transition probability of going from *i* to *i* + 1 cancer cells on a given step (*P_i,i_* _+1_) or from *i* to *i −* 1 on a given step (*P_i,i−_*_1_).

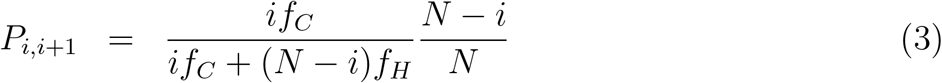

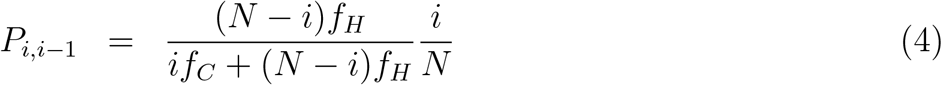

The first term in each each equation represents the probability that a cell is selected for reproduction (weighted by fitness). The second term represents the probability that a cell is selected for death. The probability of the number of cancer cells remaining the same (*P_i,i_*) is given by the following. There are two absorbing states (*P*_0,0_, *P_N,N_*).

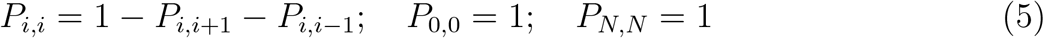

## 1 Introduction

Low dose metronomic chemotherapy (LDM) is the systematic and frequent delivery of chemotherapeutic agents at doses lower than the maximum tolerated dose paradigm (MTD) (3, 4). It is typically given at a low dose between 1/10th and 1/3rd of the maximum tolerated dose, without a long period of time between subsequent doses, hence it is also associated with higher dose densities (3). Important features of low dose, high density metronomic chemotherapy include: regular administration of chemotherapy without any interruptions using an optimized dose; preference for oral drugs; low incidence of side effects; low risk of developing resistance; lower cost. In addition, some elderly or frail patients may only be suited for lower dose chemotherapy. Residual toxicity from previous treatment may also reduce consideration for MTD chemotherapy (4). Metronomic chemotherapy regimens have been associated with lower cost of inexpensive oral drugs such as cyclophosphamide and result in fewer side-effect associated expenditures. Several phase II studies have shown promises of metronomic-like chemotherapy and its excellent safety profiles (4). The lower doses of metronomic chemotherapy regimens are now thought to not only reduce the harmful side effects of toxicity delivered to the patient but perhaps also improve anti-tumor effects (5), by killing endothelial cells in addition to its cytotoxic effect on cancer cells (6, 7) in an uninterrupted schedule for prolonged treatment periods. Metronomic chemotherapy has been shown to be effective in preclinical trials where cancer cells have developed resistance to the same chemotherapeutics (5). These LDM regimens are also suited to combination or additive strategies to new targeted and relatively non-toxic anticancer drugs recently developed.

While the advantages of LDM chemotherapy may be wide ranging with respect to toxicity, resistance, and anti-angiogenic effects (outside of the scope of our model), the goal of this article is to use an evolutionary mathematical model of cell/tumor growth with the ability to simulate chemotherapeutic scheduling to identify *growth regimes* where LDM would likely outperform MTD, and to test various scheduling protocols altering dose density and concentration. While there is no simple answer to the question of what types of tumors and growth regimes where LDM would be preferable to MTD, our results show that LDM chemotherapies with an adequate dose can outperform MTD, especially for fast growing tumors that thrive on long periods of drug-free rest with unhindered regrowth. This effect is not evident after a single cycle of chemotherapy, but is magnified after each subsequent cycle of repeated chemotherapy. In the interplay of choosing between high dose chemotherapy (MTD) or low dose, high density chemotherapy (LDM), our results show that increasing dose has diminishing returns, so the higher densities afforded by LDM regimens are an ideal tradeoff. These results may have remained hidden even in the advent of helpful theoretical regression laws like Skipper’s laws and the Norton-Simon hypothesis because these laws rely on instantaneous rates of regression, rather than the net result of the full chemotherapy cycle operating in an environment with variable growth rates. We explain how our results add to the understanding of these classic growth models and advocate the consideration of tumor growth rates when choosing chemotherapy scheduling.

### 1.1 Administration of metronomic chemotherapy

A systematic literature review of the MEDLINE, EMBASE, CENTRAL, and PubMed databases for LDM chemotherapy trials from 2000 to April 2012 performed by Lien et. al. in 2013 (4) revealed a wide variety in dose delivered and dose schedules under the terminology of metronomic chemotherapies. From the 80 studies analyzed, 107 unique treatment regimens were found (including regimens where multiple drugs were used metronomically). 38 regimens used LDM only (monotherapy n = 24, doublet LDM therapy n = 14). Of the monotherapy, the relative dose intensity (RDI: measured with respect to the maximum tolerated dose) ranged from 0.27 to 1.58 (median 1.02) and dose density (percentage of days drug is delivered) ranged from 32% to 100% (4). RDI is calculated by dividing the dose intensity (DI; the sum of the doses given each day of the chemotherapy cycle) for a chemotherapy regimen by the baseline DI value of the conventional MTD schedule. A chemotherapy may deliver a greater overall DI than the MTD (i.e. RDI *>* 1) if a lower dose is delivered more often, achieving a greater total dose over the course of the full chemotherapy cycle. The lower dose reduces toxicity, allowing for more frequent dosing, a key idea behind the metronomic schedules.

For a low dose metronomic chemotherapy, any schedule that administers a lower dose at more frequent intervals (higher dose density) could be classified as “low dose metronomic.” But, as seen above, in clinical practice the relative dose intensity delivered and the density of the scheduled are varied without clear consensus. In fact, only one monotherapy treatment regimen kept the RDI constant, balancing the lower dose with an equivalent increase in dose density. Of the remaining 23 regimens about half increased RDI (n = 12) while half decreased RDI (n = 11). It is evident that many of the quantitative details of LDM chemotherapy are unresolved including patient selection, choice of drug (or combinations of drugs and treatments), and optimal dose and treatment intervals (4). With this in mind, the goal of this manuscript is very targeted. We wish to quantify the relationship between *dose* and *dose density* delivered using the Shannon entropy index (8) as a quantitative scheduling and dosage tool. We will first briefly review the prisoner’s dilemma evolutionary game theory model of primary tumor growth that we use to carry out our computational trials (9, 10) as well as the notion of Shannon Entropy as an index to compare chemotherapeutic regimens in order to show that high entropy schedules (with an adequate dose intensity) outperform low entropy schedules.

### 1.2 The classic tumor regression laws

Benzekry et al.(11) chronicle that, despite a rise in personalized and precision medicine, currently chemotherapeutic agents are often still administered in the maximum tolerated dose paradigm. The author predicts that the forthcoming development of metronomic chemotherapy may pave the way for implementing “*computational oncology at bedside, because optimizing metronomic regimen should only be achieved thanks for modeling support*.” This prediction characterizes a growing field sometimes referred to as computational or mathematical oncology (12, 13). First, however, in order to properly understand how alternative dosing schedules like the metronomic regimens fit into the future of chemotherapy scheduling, it is important to remember the reasons that led to the advent and continued use of MTD paradigms.

#### 1.2.1 Skippers Laws

The relationship between dose and tumor cytotoxicity is linear-log (i.e. exponential decay) (14). Skipper et al. (15) were the first to develop a set of theoretical laws governing the behavior (and imply the design) of chemotherapy schedules in cancer in the late 1970s. Our understanding of the Gompertzian growth of tumors have made the application of these laws more complex, but the fundamentals of these laws still apply today (16).

In a tumor that grows exponentially (eqn. 6 and 7) with a constant exponential rate, the first law states that the tumor volume doubling time is constant over the life of the tumor (*d_t_* = log(2)*/α*),

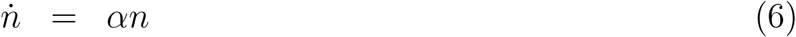

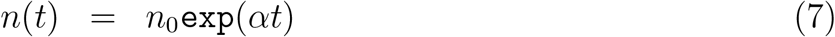

The second of Skipper’s laws is that the percentage of cells killed by a given drug dose, *D*, is constant, therefore a linear increase in dose causes a log increase in cell kill (13). As an example, a drug dose, *x*, that shrinks tumor size from 10^6^ to 10^5^ cells results in a 90% decrease of tumor population. An identical subsequent drug dose (a total dose of 2*x*) will further reduce tumor population size according to that same kill constant, to 10^4^. A third dose results in 10^3^ cells, a fourth, 10^2^, and so on. The kill law is known as a the ‘log’ kill because the constant fraction is a constant logarithmic amount. Skippers log-kill law indicates that subsequent dosing has a diminishing return; the last few remaining cells are the most difficult to eliminate. This log-linear relationship can be formulated as follows:

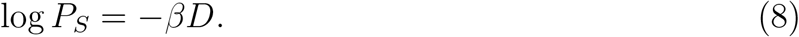

#### 1.2.2 Norton-Simon Hypothesis

One important reason the Skipper-Schabel-Wilcox model is so meaningful is that it conceptualizes the tumor growth model (e.g. exponential) and tumor regression (log-kill). Norton and Simon realized the importance of extending these observations to a Gompertzian growth model (eqn. 11). The log-kill law, a fundamentally static law does not say anything about the relationship between the fraction of cells killed and the growth rate of the tumor, only the relationship between the rate of tumor regression and the dose. In effect, Skipper’s second law assumes a constant growth rate, and therefore, a constant regression rate. In Gompertzian growth, the non-constant growth rate results in a range of log-kill rates (*β*) corresponding to the instantaneous growth rate (*γ*(*t*)). Gompertzian growth is given by the following coupled ordinary differential equations.

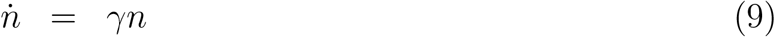

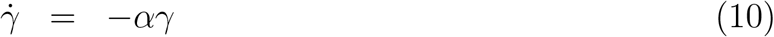

The Gompertz function reduces to the exponential function when *α* = 0. These coupled ordinary differential equations may be directly solved, as follows.

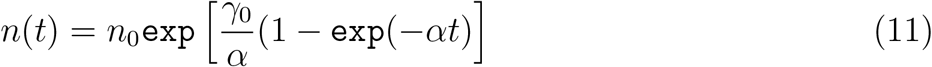

The Norton-Simon hypothesis states that tumor regression is positively (linearly) correlated with the instantaneous growth rate just before the treatment of the unperturbed tumor (17, 18). Generally, smaller tumors are associated with higher growth rates (and therefore, higher regression rates). Mathematically, the Norton-Simon Hypothesis can be formulated,

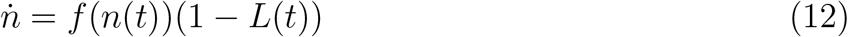

where *n*(*t*) is the growth rate model of the tumor at time *t*, *f*(*n*(*t*)) is the growth dynamics associated with the unperturbed tumor (i.e. exponential growth or Gompertzian growth), and *L*(*t*) is the loss function of cells resulting from treatment. The growth function *f*(*n*(*t*)) may be assumed to be exponential, (eqn. 7) or Gompertzian, (eqn. 9 and 10). Remembering that Skippers second law states that cell kill follows first-order kinetics, we may assume for the time being that *L*(*t*) *≡ const*., or that the rate of cell removal due to treatment is constant. In other words, each dose of chemotherapy is associated with some value of *L*. The goal is to find the optimal dose concentration and dose density that maximize the loss rate of cell kill, *L*.

### 1.3 rThe Implications of the Norton-Simon Hypothesis

Norton and Simon hypothesized that chemotherapy will only be effective in targeting cells that are in active proliferation (and as such are directly contributing the growth of the tumor in equation 12). Their model demonstrated ability to fit data preclinical experiments (19) and predict future tumor growth and regression after a few initial measurements and data from clinical trials in breast cancer (18).

The model has several key implications. First, the model predicts a higher regression for higher dose delivered. The highest dose tolerable to the patient should be chosen. Second, tumor regrowth during rest periods of chemotherapy necessitates a shorter rest period and subsequently, a shorter time of tumor regrowth. The next round of a dose dense chemotherapy will attack a smaller tumor (with higher growth rate) and lead to higher regression. Both implications give rise to the invention of the MTD paradigm to attack the tumor with the highest dose, coupled with shortest rest. These predictions were confirmed by clinical trials in which chemotherapy schedules were densified from 21 to 14 days (20). The hypothesis also predicts that tumors with an identical tumor burden may have varied responses. The growth rate of the tumor determines the response to chemotherapy. As such, early administration is important, implying a better response when the tumor is in initial stages of high growth. Similar models using the ratio of tumor volume to the host-influenced tumor carrying capacity (which corresponds inversely to the instantaneous growth rate of the tumor) has been shown to inversely predict radiotherapy response (21).

Fundamentally however, the Norton-Simon hypothesis provides no predictions for the effect of dose and dose density on regression. The Norton-Simon hypothesis (equation 12) conceals the fact that the rate of cell-kill, *L*(*t*) will be dependent on two factors: drug concentration and the number of days the drug is administered. The goal of this manuscript is to extend the classical and well-accepted predictions of Norton-Simon hypothesis from instantaneous regression rates (i.e. the derivative) to the cumulative effect (i.e. the integral) over one (or many) cycles of chemotherapy. Chemotherapy “strategies,” or schedules are quantified using the Shannon entropy (8) by their total cell reduction (TCR) over the course of the full schedule, rather than the initial regression rate (*β*). The evolutionary model we introduce in this paper is compared with regression data from murine models (see figure 3) and shown to be in good agreement.

## 2 Materials and methods

### 2.1 Chemotherapeutic agents alter the fitness landscape

It is now well established that cancer is an evolutionary and ecological process (22, 23). Studying cancer as a disease of clonal evolution has major implications on tumor progression, prevention and therapy (24, 25). The evolutionary forces at play inside the tumor such as genetic drift with heritable mutations and natural selection operating on a fitness landscape are influenced by tumor microenvironment and the interactions between competing cell types. Increased selection will influence the rates of proliferation and survival, which cause the population of cells within a tumor to progress toward more invasive, metastatic, therapeutic resistant cell types. The role of chemotheraputic agents is to kill proliferating cancer cells. This effect can be viewed as a change in fitness landscape (altering the evolutionary trajectory of the tumor), which we model as a change in the selection pressure, explained in more detail below.

In order to model these complex evolutionary forces in cancer, many theoretical biologists have used an evolutionary game theory (EGT) framework, pioneered by Nowak, to study cancer progression (see (26, 27, 28, 1, 29)). Evolutionary game theory provides a quantitative framework for analyzing contests (competition) between various species in a population (via the association of strategies with birth/death rates and relative sub-clonal populations) and provides mathematical tools to predict the prevalence of each species over time based on the strategies (30, 31, 28, 32). More specifically, the framework of EGT allows the modeler to track the relative frequencies of competing subpopulations with different traits within a bigger population by defining mutual payoffs among pairs within the group. From this, one can then define a fitness landscape over which the subpopulations evolve.

### 2.2 The model

The model presented in (9, 10) and used in this paper to carry out our computational trials is a framework of primary tumor growth used to test the effect of various chemotherapuetic regimens, including MTD and LDM. The model is a stochastic Moran (finite-population birth-death) process (33) that drives tumor growth, with heritable mutations (34) operating over a fitness landscape so that natural selection can play out over many cell division timescales (described in more detail in (9, 10)). The birth-death replacement process is based on a fitness landscape function defined in terms of stochastic interactions with payoffs determined by the prisoner’s dilemma game (1, 2). This game incorporates two general classes of cells: healthy (the cooperators) and cancerous (the defectors) (35, 36). During tumor progression, each cell is binned into one of two fitness levels, corresponding to their proliferative potential: healthy (low fitness) and cancer (high fitness). In our model, we can think of a cancer cell as a formerly cooperating healthy cell that has defected and begins to compete against the surrounding population of healthy cells for resources and reproductive prowess. The model demonstrates several simulated emergent ‘cancer-like’ features: Gompertzian tumor growth driven by heterogeneity (37, 38, 39), the log-kill law which (linearly) relates therapeutic dose density to the (log) probability of cancer cell survival, and the Norton-Simon hypothesis which (linearly) relates tumor regression rates to tumor growth rates, and intratumor molecular heterogeneity as a driver of tumor growth (10).

Others have presented mathematical models to study evolutionary dynamics of tumor response to targeted therapy (40) in either combination or sequential therapy (41, 42), and optimal drug dosing schedules to prevent or delay the emergence of resistance or optimize tumor response (43, 44, 45). We are interested testing “strategies,” or drug schedules that control the number of cancer cells, *i*, in a population of *N* cells comprising the simulated tissue region. (Note: the size of the tumor population, *i*, is variable and changing according to the fitness landscape, detailed in equations 1 through 5. The carrying capacity, *N*, is a parameter in the model, but all plots shown here are normalized by *N*, so the proportion of cancer cells, *i/N*, is used to track tumor growth, without loss of generality.) The model presented here uses a parameter, *w*, to control the effect of selection pressure. A value of *w* = 0 corresponds to neutral drift (no selection) and a value of *w* = 1 corresponds to strong selection. We break *w* into two separate parameters, *w_H_*, the selection pressure on the healthy population, and *w_C_*, the selection pressure on the cancer population (see figure 1a). Each dose of chemotherapy is associated with a dose concentration, *c*, which alters the selection pressure as indicated in figure 1a. Here, we assume drug concentration will be measured as as a fraction of the conventional maximum tolerated dose (MTD) dosages, hence 0 *≤ c ≤* 1 (see Figure 1b). As *c* increases, the selection pressure is altered in favor of the healthy cells (*w_H_ > w*) and to the detriment of cancer cells (*w_C_ < w*) as shown in Figure 1a. Before therapy, the fitness landscape of an untreated tumor is that of a prisoner’s dilemma (Figure 1c), where the fitness of the cancer subpopulation is greater than the healthy population for the entire range of cancer proportion, *i/N*. The change in fitness landscape for a moderate value of *c* (*c* = 0.4) is shown in Figure 1d, which gives the healthy cell population a fitness advantage over the cancer population. The advantage is lessened as the tumor size (*i/N*) increases (which contributes to the emergence of the Norton-Simon model in Figure 2, explained in detail below). For a strong dose of therapy (such as *c* = 0.8, shown in Figure 1e), the effect on the fitness landscape is exaggerated. Thus, a higher dose leads to a higher kill rate of cancer cells.

**Figure 1:**
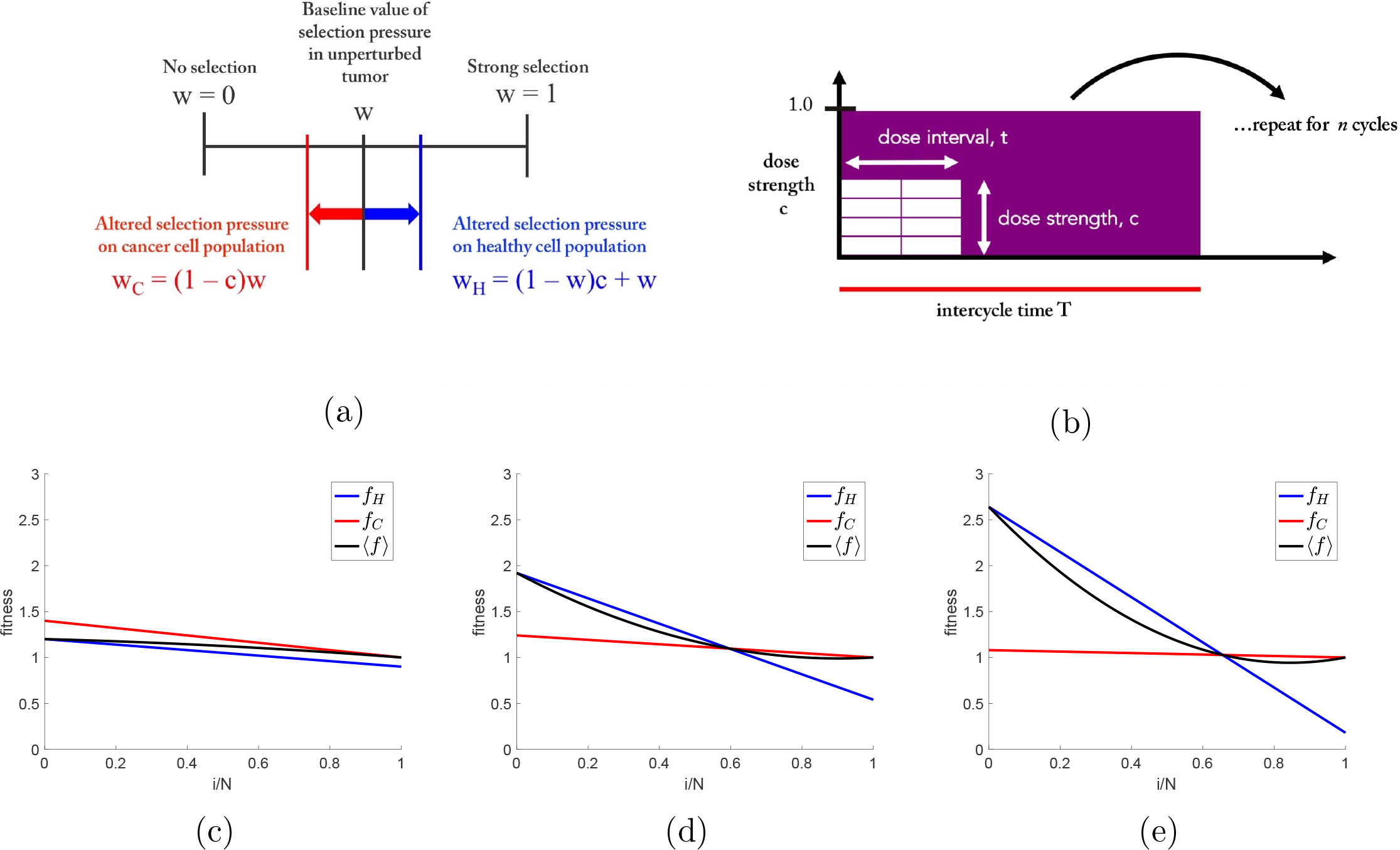
**Chemotherapy is a selective agent that alters the fitness landscape of cells** — (a) The dose strength parameter, *c*, (0 *≤ c ≤* 1), alters the selection pressure parameter, *w*, (0 *≤ w ≤* 1), in favor of the healthy cell population (*w_H_ > w*) and to the disadvantage of the cancer cell population (*w_C_ < w*). (b) Total dose density delivered in the one chemotherapeutic cycle, D, is the product of the dose strength (*c*, 0 *≤ c ≤* 1) and dose interval (*d*, 0 *≤ d ≤* 1) such that *D* = *ct* (eqn. 13 (0 *≤ D ≤* 1). (c,d,e) Plots showing the fitness of the healthy cell subpopulation (*f_H_*, blue) and the cancer cell subpopulation (*f_C_*, red) for no therapy, low dose therapy, and high dose therapy.

In literature, two models have been proposed to model loss functions due to a drug: 1) non-cycle specific (where the loss function is linear with tumor size) (46) and 2) cycle-specific (where loss function is linear with tumor growth rate) (15, 18). Cycle-specific drugs are considered here, and thus a model of regression that is linear with tumor growth rate is chosen. The loss function of the Norton Simon hypothesis in equation 12 shows an example of cycle-specific drug modeling.

Average instantaneous growth rates of stochastic Moran process models (see equations 1 through 5) are proportional to fitness: *γ ∝ w*(*f_i_* − 〈*f*〉), where *f_i_* is the fitness of cell type *i*, *(f)* is the average fitness of total population, and *w* is the selection pressure parameter, as explained in the Key Equations section, and shown in figures 1c, 1d, and 1e. This proportionality between growth (*γ*) and selection pressure (*w*) indicates that varying *w* linearly with dose concentration *c* (shown in figure 1a) is directly comparable to the previous cycle-specific drug models. This linearity is confirmed by plotting instantaneous regression rates throughout the life of a tumor in figure 2. Identical, continuous chemotherapy is administered at different time points in the life of the tumor, corresponding to different instantaneous growth rates (figure 2a). A linear relationship between the instantaneous growth rate (*γ*) and instantaneous regression rate (*β*) (figure 2b) emerges from the model, comparable to the predictions of the Norton-Simon hypothesis.

**Figure 2:**
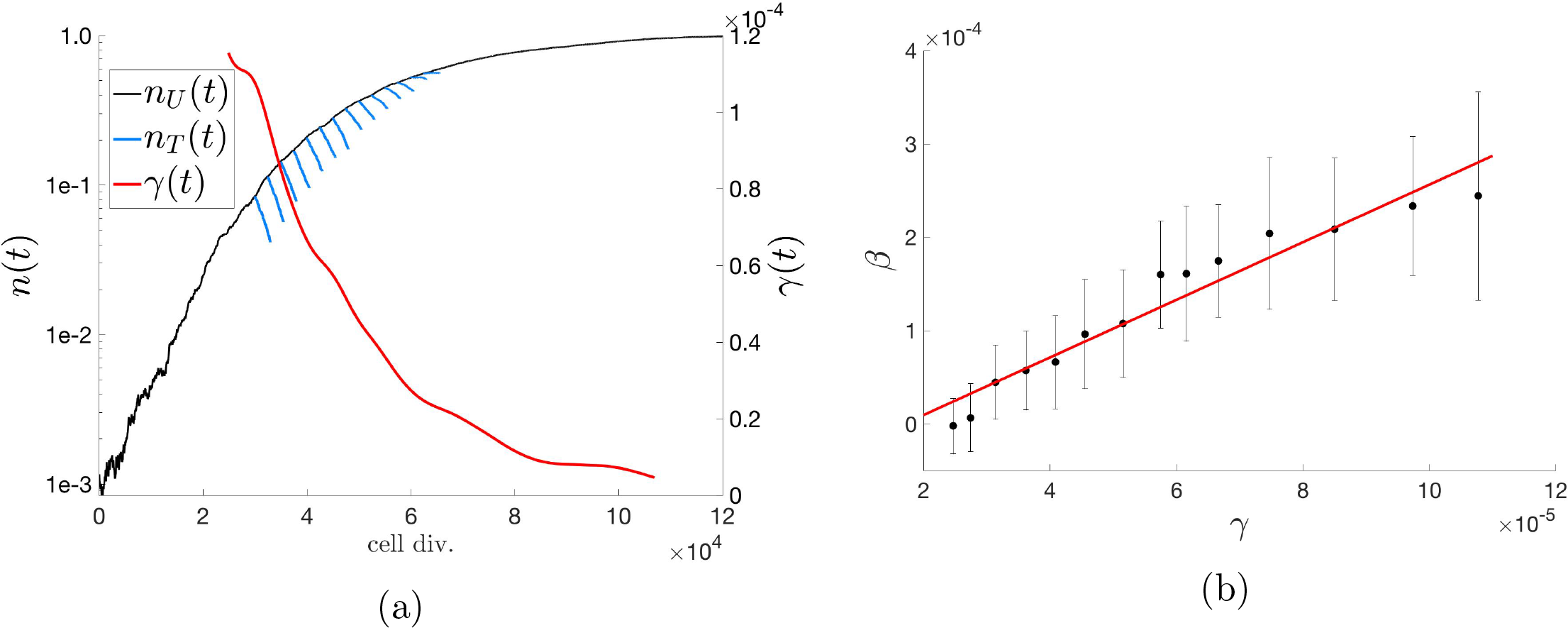
**Classical Tumor Regression Laws** — (a) The Norton-Simon hypothesis states that tumor regression is proportional to the growth rate of an unperturbed tumor of that size. Unperturbed tumor growth, *n_U_* (*t*) (black) in a representative population of *N* = 10^3^ cells, and growth rate, *γ*(*t*) (red) is shown. Therapy is administered at various timepoints in the growth of the tumor and then regression, *n_T_* (*t*), is plotted (blue). Rate of regression, *β*, is the best-fit slope on the log-plot. (b) The average regression rate was calculated for 25 stochastic simulations, and plotted as a function of *γ* at the time of therapy with error bars indicating the standard deviation of values. A linear best fit (predicted to be linear by the Norton-Simon hypothesis) is calculated to be *β*(*t*) = 3.0865*γ* + 5.2359*e*05.

A separate justification of the linear model of the effect of drug concentration on selection pressure is shown in figure 3. A best-fit was performed to find the optimal parameters of *w* and *c* to fit data reproduced from mouse models quantifying inter-mouse and intra-mouse variability and response to 5-Fluorouracil (5-FU) in two treatment groups: 50mg/kg (figure 3 left panel) and 100mg/kg (figure 3 right panel) (47). Tumor size measurements were taken from visibility until a mouse tumor reached 3 to 4 mm in size, and drug treatment was administered weekly until 1cm in size. The prisoner’s dilemma model (black solid lines) appears to accurately capture both the growth dynamics (solid black circles) and the treatment dynamics (black x’s). The dashed lines are Gompertzian best-fit functions of the unperturbed pre-treament data (black circles), showing good aggreement with the prisoner’s dilemma (black solid lines). Previously, we have reported the model’s success in capturing current unperturbed growth models (i.e. Gompertzian growth) as an emergent phenomena of this evolutionary model (10).

**Figure 3:**
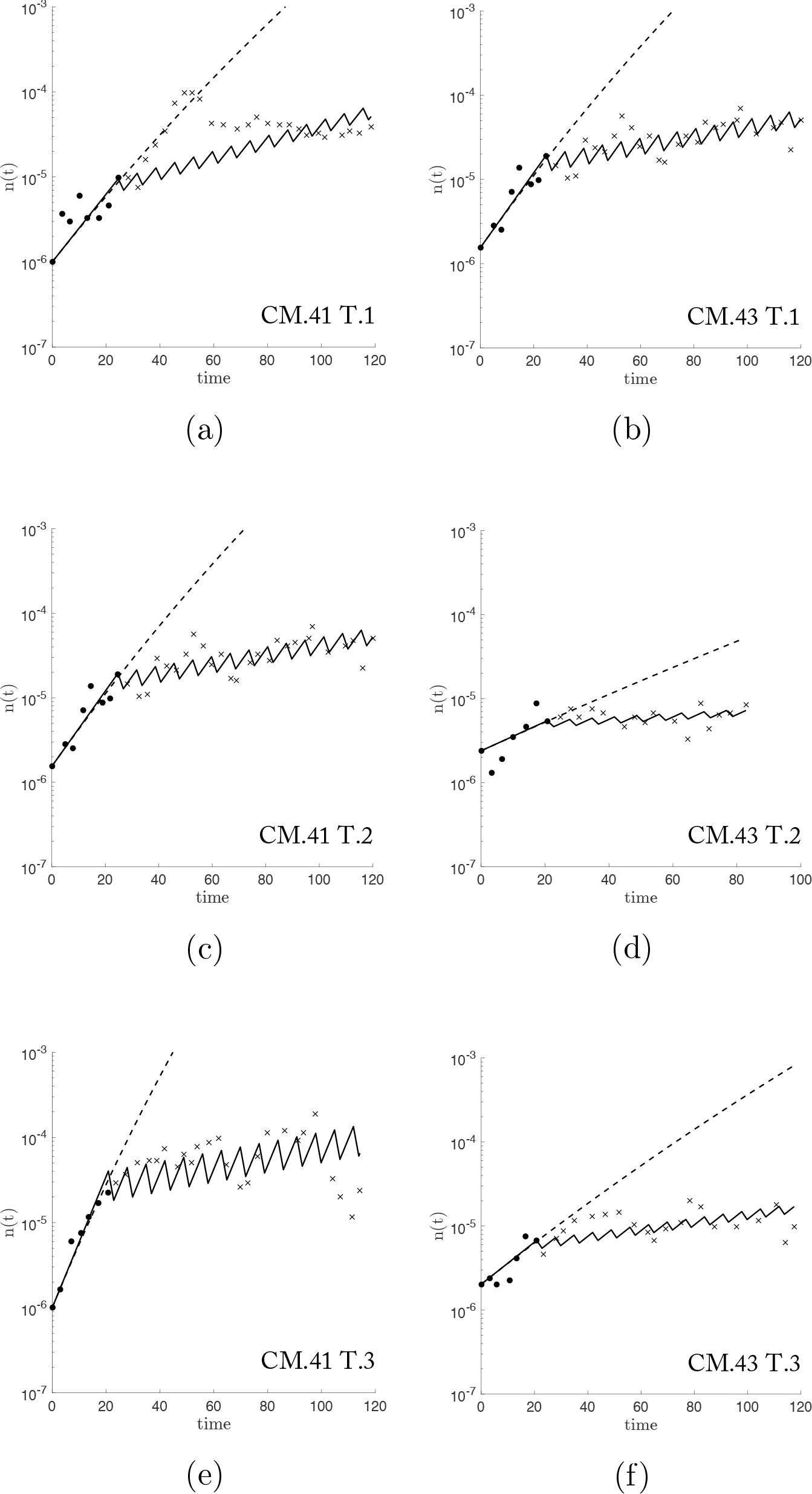
**Response of murine tumors to 5-Fluorouracil (5-FU) treatment with model best-fit** — Data (reproduced from (47)) from two treated mice: CM.41 (left panel) and CM.43 (right panel), receiving doses of 50mg/kg or 100mg/kg, respectively, on 2 days out of 7 (biweekly). Biweekly measurements of tumor volume are recorded for untreated (black circles) until 3-4mm in size and treated volumes (black x’s) are measured until tumor reaches 1cm size. A Gompertzian function is best-fit (dashed line) and the Prisoner’s dilemma model is fit using *w* and *c* as parameters (solid line). The model fit performs well for the wide range of tumor growth rates found in six tumors (*w* = [0.18, 0.08, 0.21, 0.08, 0.35, 0.12] and *c* = [0.30, 0.49, 0.34, 0.34, 0.36, 0.32] left to right, top to bottom, respectively). Note: tumor in (a) shows a time delay from start of treatment to response to therapy which our model does not address.

### 2.3 Dose concentration versus dose density

Despite a growing trend toward personalized and precision medicine, treatment goals have shifted from complete cure to an optimization of long term management of the disease; rather than trying to find the silver bullet, we might utilize the advances in mathematical models to optimize existing therapeutic options (11, 48). For this reason, we have decided to test the merit of various chemotherapeutic regimens by comparing the total tumor cell reduction (TCR). Presumably, a therapy regimen with a higher value of TCR will provide a greater level of tumor control, a longer time to relapse, and better prognosis.

A drug dose, *D*, (equation 13) is generally measured in units of mg/m^2^/week (here, average body surface area assumed to be 1.8m^2^). Yet, dose *D* consists of two components: dose concentration (parameter *c* in our model) and dose time factor (parameter *t* in our model). The time factor, called the dose density when normalized by the intercycle time, represents the percentage of days a dose is administered. In order to compare the importance of each term on tumor cell reduction, we hold one term constant and vary the other in Figure 4.

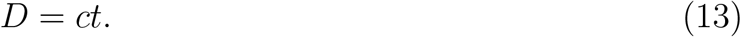

**Figure 4:**
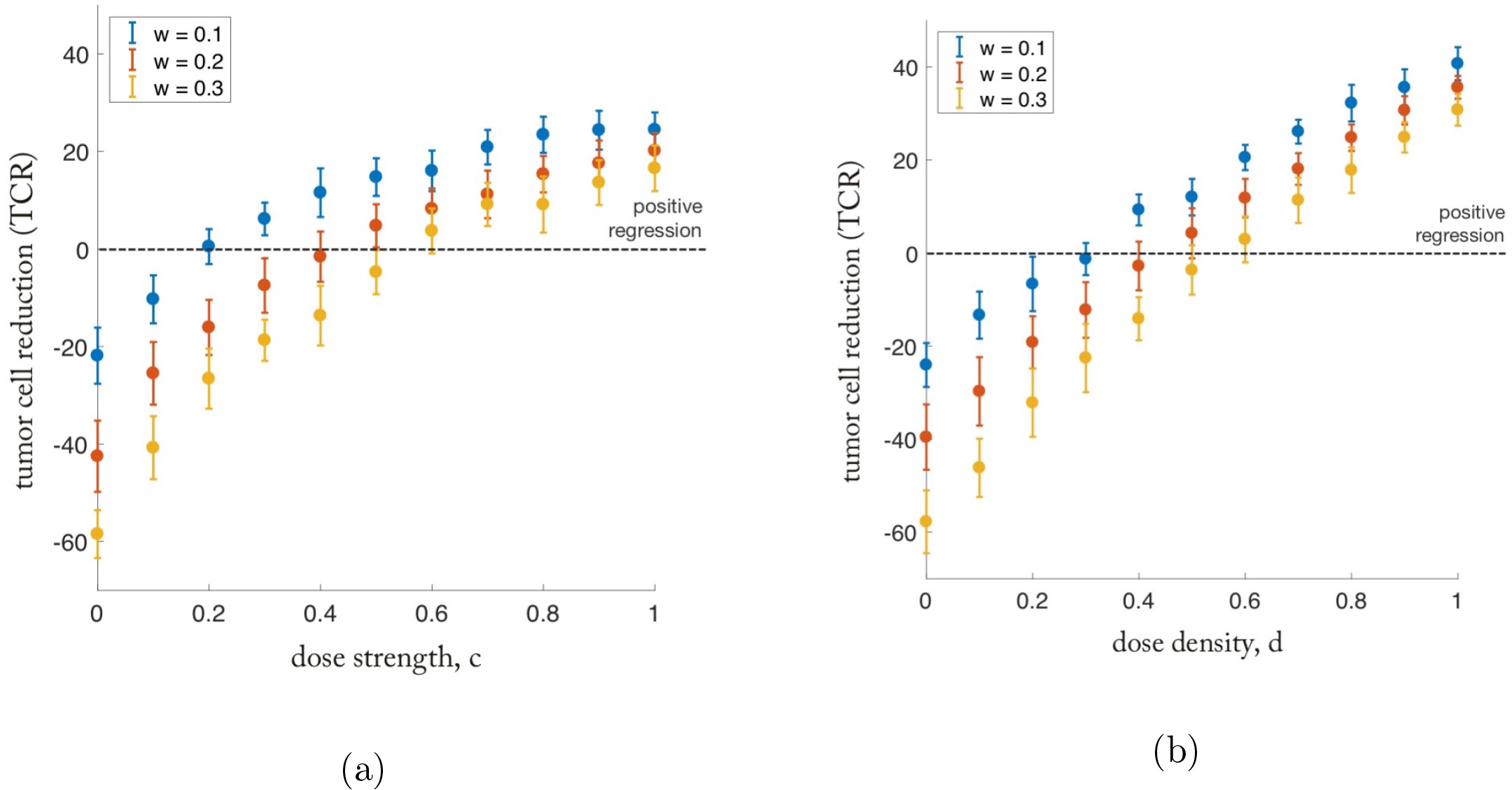
**Diminishing returns of dose escalation compared to linear relationship of dose density** — (a) Dose Escalation: The percent regression of a tumor for a range of dose strength (constant dose interval: *t* = 10 days, *T* = 14 days) are shown for a range of selection pressure: *w* = 0.1 (blue), *w* = 0.2 (red), and *w* = 0.3 (yellow). For each subsequent increase in dose strength, the dose escalation approach to chemotherapy shows diminishing returns in percent tumor regression. (b) Dose Density: The percent regression of a tumor for a range of dose interval (constant dose strength: *c* = 1.0) are shown for a range of selection pressure: *w* = 0.1 (blue), *w* = 0.2 (red), and *w* = 0.3 (yellow). Dose density shows a linear relationship between densifying chemotherapy and percent tumor regression.

Clearly seen in Figure 4a, there is a diminishing return on increasing the dose strength of a given chemotherapy regimen. Although there is a positive relationship (an increase in dose leads to a higher regression) that relationship lessens as the dose is increased further. However, in Figure 4b, the relationship between dose density and regression is linear, showing no signs of diminishing returns of increasing density.

The point has an important subtlety: the dose cannot be continually lowered in favor of density. The dose must be sufficient to overcome the growth rate of the tumor; some doses are not adequate for tumor regression regardless of the density. This is seen for values below the dotted line in Figure 4a and 4b.

## 3 Results

### 3.1 Quantifying chemotherapeutic strategies via entropy metric

In clinical practice today, there are three common chemotherapy regimens in use considered here: Maximum Tolerated Dose (MTD), Low Dose Metronomic weekly (LDMw) and Low Dose Metronomic daily (LDMd). These three chemotherapy strategies are shown in Figure 5, left. Each regimen consists of identical cycles that are repeated until the tumor is eradicated. The MTD (left, top) regimen delivers the maximum dose on a single day, repeated once every 2 weeks. The LDMw (left, middle) regimen lowers the dose, but doubles the dose density from 1 to 2 days out of 14. The LDMd (left, bottom) regimen has the highest density (there is a dose administered on 100% of the days), but the lowest dose.

**Figure 5:**
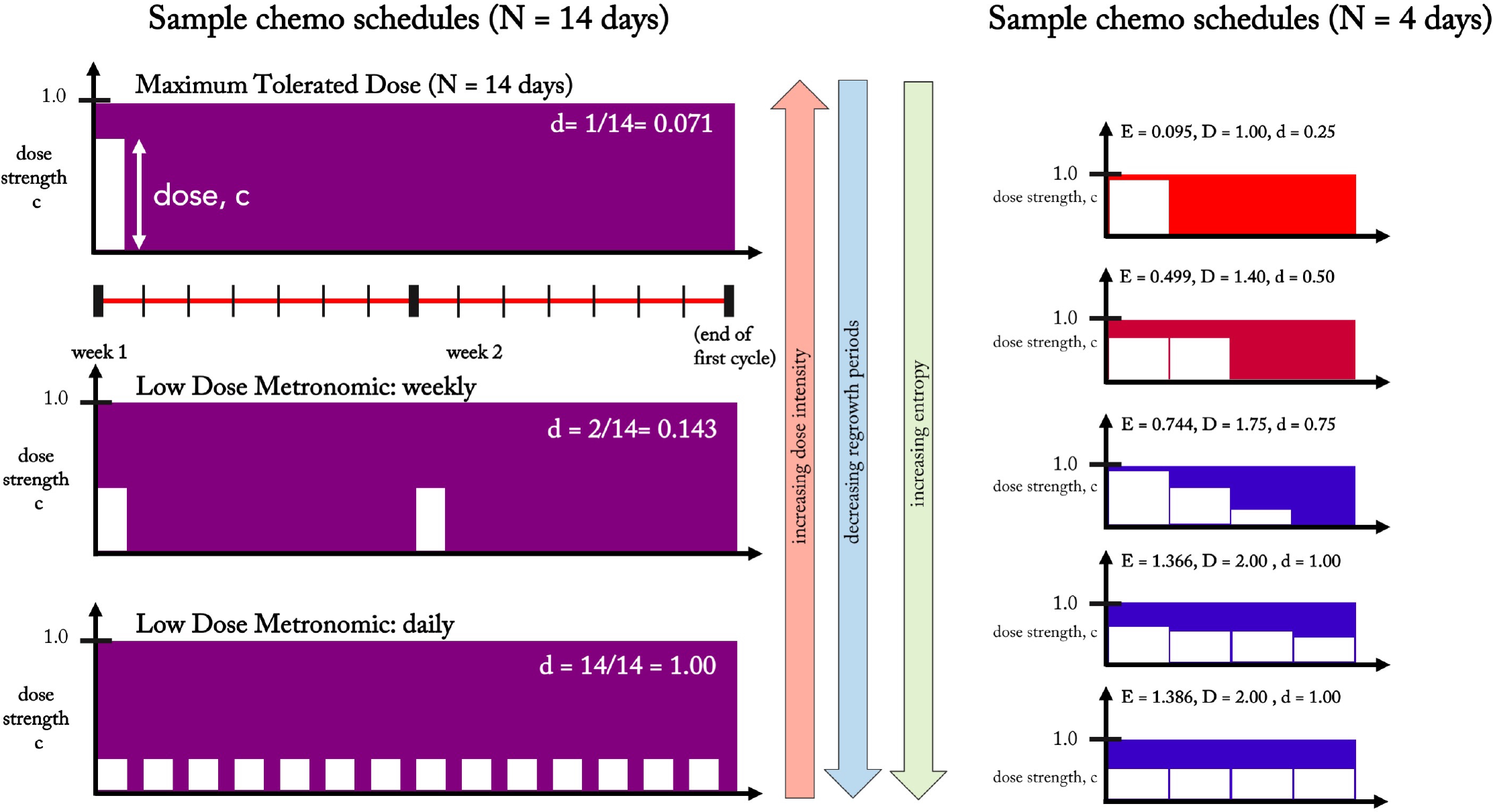
**Shannon entropy as an index to compare treatment strategies** — **Left:** 3 common chemotherapy schedules are shown for one cycle (*N* = 14 days). Maximum Tolerated Dose (left, top) is a high dose (administered once at the beginning of every 2 week cycle) and low dose density (*d* = 0.071, see equation 16) regimen. Low Dose Metronomic Weekly (left, middle) is a lower dose, higher density (*d* = 0.143) regimen, while Low Dose Metronomic Daily is the lowest dose, highest density (*d* = 1.00). **Right:** Similarly, chemotherapy regimens can be simulated for a range of dose, density, and entropy values. Pictured from top to bottom are a range of representative regimens from low entropy (i.e. high dose, low density) to high entropy (i.e. low dose, high density) for a cycle of *N* = 4 days. On each *i*th day, treatment of dose *c_i_* is administered. The treatment strategy’s Shannon Entropy, *E*, is calculated according to equation 14 and the total dose delivered is calculated according to equation 15. All treatment strategies are front loaded (monotonically decreasing) regimens. It should be noted that LDM-like regimens correspond to a high entropy value (bottom, left and right).

There are hundreds of such choices of chemotherapy regimens when considering varying doses across many days or weeks (Figure 5, right), each varying the total dose delivered, *D*, and the density, *d*. We propose using a Shannon Entropy index, *E*, of a given chemotherapy schedule as a measure that can quantify and synthesize information of both the dose on a given day and the distribution of unique, daily doses across the entire chemotherapy regimen into a single metric. The entropy is calculated as follows, where *c_i_* is the dose strength (often simply referred to as ‘dose’) on day *i*.

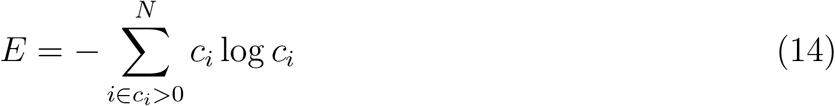

The assumption in equation 13 that an identical dose is delivered every day can be relaxed, and the total dose delivered is found by summing the dose on each *i*th day (*c_i_*) multiplied by the length of the dose in days (*t_i_*). We assume that the smallest resolution of discrete times between doses, *t_i_* is a single day, or *t_i_* = 1 for all *i*. *N* is the number of days between cycles, also known as the intercycle time.

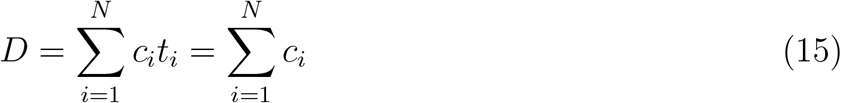

The dose density of a regimen can be found by summing the number of days where a non-zero dose is delivered, and dividing by the intercycle time in days, *N*. Thus, the density will be a non-dimensionalized parameter such that (0 *≤ d ≤* 1).

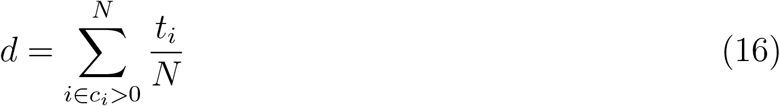

The Shannon entropy metric, *E*, is an ideal metric for comparing chemotherapy regimens because it separates the existing cases already in clinical practice today: MTD (low entropy, characterized by high doses with long periods of rest), metronomic regimens (high entropy, characterized by low doses with short or no periods of rest), as well as any arbitrary strategy of varied doses administered in a cycle of arbitrary length of days. All of the simulated therapy regimens were assumed to be frontloaded (non-increasing, with the highest dose on day 1 and equal or lower subsequent doses). Backloaded regimens give similar but slightly disadvantageous results, because backloaded regimens often start with a period of rest, giving the tumor time to grow to a larger tumor, which is associated with a lower growth rate (and therefore lower regression).

### 3.2 LDM versus MTD chemotherapies

Computational simulations of 1000 unique chemotherapy schedules were run with identical initial conditions (*N* = 1*e*6 cells; *i/N* = 1*e*3). Mean values of tumor cell regression percentage for 50 simulations were calculated and plotted in a pictorial histogram according to regression percentage (Figure 6). Both slow growing tumors (*w* = 0.1) and fast growing tumors (*w* = 0.2) were simulated.

**Figure 6:**
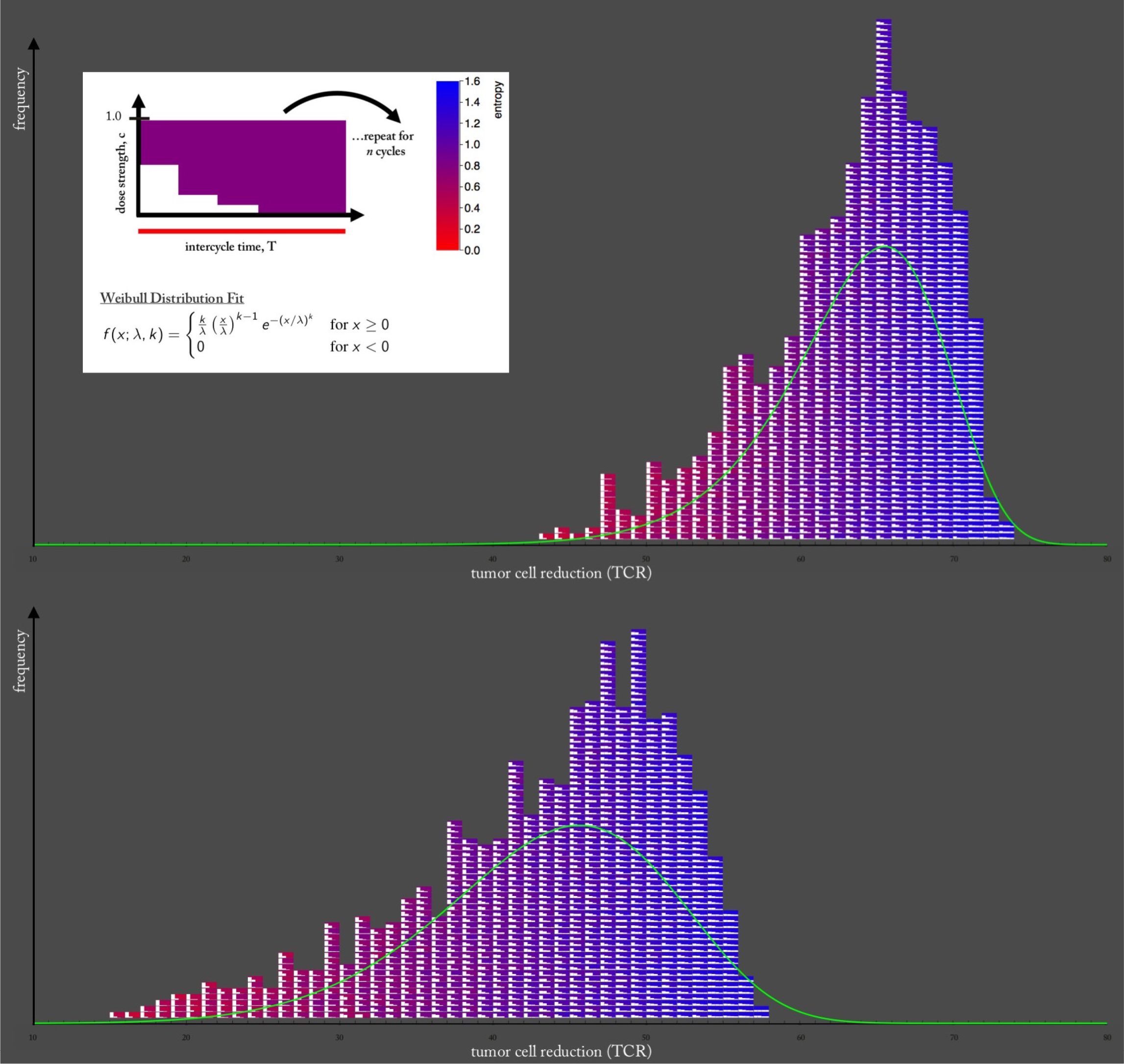
**High entropy, LDM-like chemotherapies outperform low entropy MTD-like chemotherapies** — Two pictorial histograms are plotted where each block (color-coded from red: low entropy to blue: high entropy) represents a chemotherapy regimen. The top histogram is for a slow-growing tumor (*w* = 0.1) and the bottom histogram is faster growth (*w* = 0.2). All regimens are equivalent total dose (*D* = 0.3), monotonically decreasing, and are repeated for 8 cycles of chemotherapy and the tumor cell reduction (TCR) is recorded. The dose density, *d*, and dose concentration, *c_i_*, are varied between regimens. The histogram clearly shows a color-shift from red toward blue for low TCR, ineffective therapies toward high TCR, effective therapies. High entropy (blue) therapies outperform low entropy therapies. The data was fit to a Weibull distribution (shown in upper left panel; top: *k* = 14.251, *λ* = 65.882, bottom: *k* = 6.647, *λ* = 46.758), overlaid in green.

Each block represents a chemotherapy regimen, which has an associated Shannon entropy index (eqn. 14). The background color of the blocks of the chemotherapy regimens are color-coded from red (low entropy) to blue (high entropy). The smaller white squares within each block indicate the strength of the therapy dose for each day (*c_i_*). Pictured are 1000 combinations of *N* = 4 day chemotherapy schedules, but similar trends are seen for chemotherapy schedules of longer length of days. All regimens are equivalent total dose (*D* = 0.3), non-increasing, and are repeated for 8 cycles of chemotherapy and the tumor cell regression (TCR) is recorded. The histograms clearly show a color-shift from red toward blue for low TCR toward high TCR. This indicates that high entropy (blue) therapies outperform low entropy therapies and consistently lead to higher tumor cell reduction. These high entropy regimens are low dose, more dose-dense chemotherapies, characteristic of LDM chemotherapy.

In Figure 7, the analysis is repeated for varied tumor growth rates (i.e. varied selection pressure) for *w* = 0.1, (blue) *w* = 0.2, (red) and *w* = 0.3 (yellow). The difference in reduction is shown for 1 cycle, 8 cycles, and 16 cycles. Fast growing tumors have a high slope on a least-squares linear fit approximation of the entropy-TCR plot, which means that high entropy therapies (LDM) are more effective for fast growing tumors than for slow growing tumors. By contrast, slow growing tumors have a lower slope on the entropy-regression plot, which means that all regimens have relatively similar performance outcomes. Fast growing tumors, therefore, have a higher likelihood of benefiting from a more LDM-like chemotherapy, provided the dose is adequate to lead to tumor regression.

**Figure 7:**
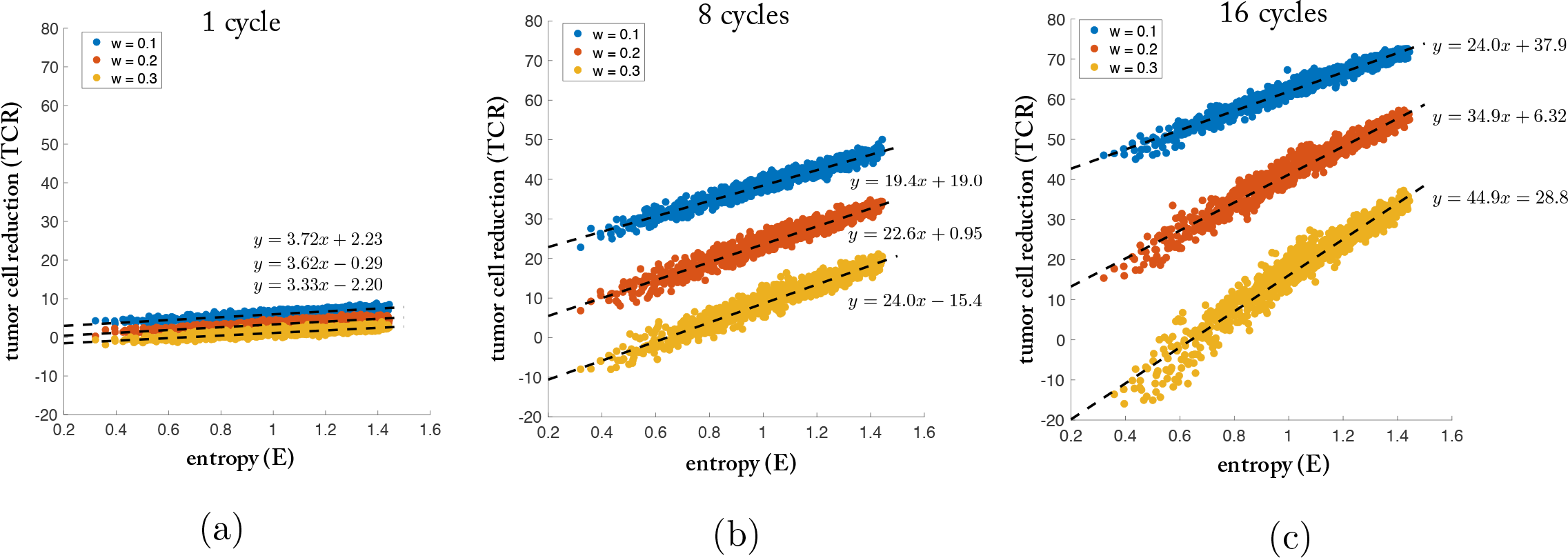
**High entropy strategies lead to an increase in tumor regression** — The relationship between tumor cell reduction (TCR) and entropy (H) is shown for a single cycle of chemotherapy (a), 8 cycles (b), and 16 cycles (c). The simulations (averages of 25 stochastic simulations for total dose delivered *D* = 0.3) are repeated for slow (*w* = 0.1, blue), medium (*w* = 0.2, red), and fast growing tumors (*w* = 0.3, yellow). The low slope value in (a) indicates negligible advantage of high entropy strategies after only a single cycle. After many cycles, the advantage of high entropy strategies is apparent (b,c). Also note that the slope associated with faster growing tumors (yellow) is higher than those of slower growing tumors (blue). This indicates that at high entropies, TCR for the fast growing tumors is closer to those for slow growing tumors, as compared with low entropies.

The effect is almost negligible after a single cycle (Figure 7a). The appeal of the implication of Norton-Simon toward an MTD approach to chemotherapy lies in the high initial response of tumors to a high dose. The metronomic chemotherapies take more cycles to overtake the initial quick response of the MTD, but after the 8 cycles (Fig. 7b) and 16 cycles (Fig. 7c), the cumulative effect is evident and metronomic chemotherapies outperform MTD therapies. For each growth rate (i.e. selection pressure), there is a corresponding optimal chemotherapy schedule. In each case, the optimal solution corresponds to the highest entropy (which corresponds to the low-dose metronomic chemotherapy schedule).

## 4 Discussion

We use a stochastic Moran process model coupled with a prisoner’s dilemma evolutionary game (cellular interactions) to contrast LDM and MTD chemotherapies with respect to their effect on tumor growth. The Shannon entropy was identified as a useful metric to compare chemotherapy strategies. The metric is useful in quantifying LDM strategies (which correspond to high entropy values), MTD strategies (low entropy), as well as novel strategies with intermediate entropy values.

Our results show that high dose chemotherapy strategies outperform low dose, although there are some subtleties associated with the growth rates of the tumors. Dosing consists of a product of concentration and density and our results show that an increase in density is more effective than the same percentage increase in concentration. In other words, higher dose concentrations shown diminishing returns. The effectiveness of density in leading to a higher tumor cell reduction allows the LDM chemotherapies (which are more dose dense) to outperform MTD strategies. This effect is magnified for fast growing tumors that thrive on long periods of unhindered growth without chemotherapy drugs present. This effect is not evident after a single cycle of chemotherapy, but is magnified after each subsequent cycle of repeated chemotherapy. We could ask if there is any evidence of this effect in the literature on clinical trials already performed. We first point to a paper comparing different chemotherapeutic schedules for prostate tumors (49) (relatively slow-growth rates). In this phase 3 study, docetaxel dosing given every three weeks was compared to dosing every week. The mean survival was only slightly higher for the first group (three weeks) compared with the second (weekly), showing no obvious benefit to a low-dose high density treatment. By contrast, a phase 2 trial for small cell lung cancer (SCLC) (50) was performed, a tumor with typically higher growth rates than prostate tumors. For this group, the drug topotecan was administered on a higher dose weekly basis with disappointing results, pointing out the advantages of the LDM therapies for this fast-growing tumor type.

Thus our model points to the benefits of choosing dosing strategies based on tumor growth rates, something not currently done in medical practice. The concept of choosing dosing schedules based on tumor growth rates could well be a fruitful avenue to test further in clinical trials focused on this question. Others have attempted to estimate prospective patient-specific tumor growth rates to make clinical decisions about treatment scheduling and fractionization, using measurements at diagnosis and first day of treatment (51, 21). Furthermore, the promise of LDM chemotherapy on mitigating the risk of resistance (5) and metastasis (11) could be a separate line of future investigation.

## Notes

**Disclosure of Potential Conflicts of Interest:** The authors declare no potential conflicts of interest.

**Financial Support:** West is supported by Award Number DGE 1045595 from the National Science Foundation.

